# Scanning sample-specific miRNA regulation from bulk and single-cell RNA-sequencing data

**DOI:** 10.1101/2023.08.21.554111

**Authors:** Junpeng Zhang, Lin Liu, Xuemei Wei, Chunwen Zhao, Yanbi Luo, Jiuyong Li, Thuc Duy Le

**Author notes:** Correspondence may also be addressed to Thuc Duy Le.

## Abstract

RNA-sequencing technology provides an effective tool for understanding miRNA regulation in complex human diseases, including cancers. A large number of computational methods have been developed to make use of bulk and single-cell RNA-sequencing data to identify miRNA regulations at the resolution of multiple samples (i.e. group of cells or tissues). However, due to the heterogeneity of individual samples, there is a strong need to infer miRNA regulation specific to individual samples to uncover miRNA regulation at single-sample resolution level. Here, we develop a framework, Scan, for scanning **<u>s</u>**ample-spe**<u>c</u>**ific miRNA regul**<u>a</u>**tio**<u>n</u>**. Since a single network inference method or strategy cannot perform well for all types of new data, Scan incorporates 27 network inference methods and two strategies to infer tissue-specific or cell-specific miRNA regulation from bulk or single-cell RNA-sequencing data. Results on bulk and single-cell RNA-sequencing data demonstrate the effectiveness of Scan in inferring sample-specific miRNA regulation. Moreover, we have found that incorporating priori information of miRNA targets can improve the accuracy of miRNA target prediction. In addition, Scan can contribute to the clustering cells/tissues and construction of cell/tissue correlation networks. Finally, the comparison results have shown that the performance of network inference methods is likely to be data-specific, and selecting optimal network inference methods is required for more accurate prediction of miRNA targets. We have made Scan freely available to the public to help infer sample-specific miRNA regulation for new data, benchmark new network inference methods and deepen the understanding of miRNA regulation at the resolution of individual samples.

## Introduction

As a type of regulatory non-coding RNA (ncRNA), microRNA (miRNA, 19-25 nt in length) can modulate the expression levels of thousands of coding RNAs, thereby playing a key role in biological processes, signaling pathways and pathological processes of human cancers^1–3^. Due to the importance of miRNAs in biological system, they have the potential to be biomarkers of human cancers, e.g. breast cancer^4^, leukemia^5^ and prostate cancer^6^ in clinical applications. In addition to the miRNA-directed regulation which results in inhibition or degradation of messenger RNAs (mRNAs), miRNAs can also act as mediators of the crosstalk between other ncRNAs (e.g. long non-coding RNAs, circular RNAs and pseudogenes) and mRNAs^7,8^. The miRNA-mediated or miRNA-indirect regulation is also known as competing endogenous RNA (ceRNA) regulation, and has been demonstrated to be involved in the regulatory mechanism of human cancers^9–11^.

There are multiple types of biological networks, including gene regulatory network within a biological sample (cell or tissue). Among the different types of biological networks, gene regulatory network is regarded as an important feature of biological samples, and gives rise to the unique characteristics of each biological sample^12^. In the gene regulation field, miRNA regulation has attracted broad attentions on account of its potential clinical translation. Therefore, inferring and characterizing miRNA regulation is a central problem for revealing miRNA regulatory mechanisms in systems biology. Just like dynamic biological system in a biological sample (cell or tissue), miRNA regulation also tends to be dynamic. For example, by using bulk and single-cell RNA-sequencing data, existing studies^13–17^ have found that the miRNA regulatory networks are condition-specific, tissue-specific and cell-specific. Moreover, it is widely known that biological samples are characterized by heterogeneity, and are shown to be unique. Thus, to understand the miRNA regulation specific to biological samples (cancer tissues or cells in particular), it is essential to investigate sample-specific miRNA regulation (i.e. the miRNA regulatory network for a specific sample).

At the single-sample level, previous methods including CeSpGRN^12^, CSN^18^, DEVC-net^19^, EdgeBiomarker^20^, SSN^21^, LIONESS^22^, c-CSN^23^ and LocCSN^24^ have been presented to identify sample-specific gene regulation (one gene regulatory network for one sample), and the identified gene regulatory networks with these methods are undirected networks. To infer causal regulatory networks (represented as directed networks) specific to individual samples, an inference method^25^ using drift-diffusion processes is also proposed. However, the above computational methods^12,18–25^ are designed for exploring sample-specific transcriptional regulation rather than sample-specific miRNA regulation. To study miRNA regulation at single-cell level, CSmiR^16^ is introduced to infer cell-specific miRNA regulation from singlelJcell miRNAlJmRNA colJsequencing data (where each single cell is considered as a sample). As stated in CSmiR, for singlelJcell miRNAlJmRNA colJsequencing data with less than 100 single-cells, CSmiR needs to interpolate pseudo-cells by using a re-sampling technique. However, the original single-cell miRNA-mRNA co-sequencing data itself contains technical noise (e.g. high dropout rate), thus as a result, the enlarged data by adding pseudo-cells may aggravate the technical noise.

To model the dynamic regulatory processes of miRNAs at the single-sample level, in this work, we propose a **<u>S</u>**ample-spe**<u>c</u>**ific miRNA regul**<u>a</u>**tio**<u>n</u>** (Scan) framework to scan sample-specific miRNA regulation from bulk and single-cell RNA-sequencing data. Due to the fact that that a single network inference method or strategy does not always perform the best in all types of new data, Scan incorporates 27 network inference methods and two strategies to infer tissue-specific or cell-specific miRNA regulation from bulk or single-cell RNA-sequencing data. Given bulk or single-cell RNA-sequencing data with or without priori information of miRNA-mRNA interactions, 27 well-used network inference methods in Scan can be used to calculate the strength between miRNAs and mRNAs. Based on the strength between miRNAs and mRNAs, Scan adapts two strategies: statistical perturbation^21^ and linear interpolation^22^ to infer sample-specific miRNA regulatory networks.

By applying Scan to two RNA-sequencing datasets, a bulk dataset from breast cancer tissues, and a single-cell dataset from chronic myelogenous leukemia cells, we demonstrate the effectiveness of Scan in terms of accuracy and efficiency of miRNA target prediction. Moreover, the performance comparison between with using and without using priori information indicates that the priori information of miRNA targets can improve the accuracy of miRNA target prediction. We also reveal that Scan can help to cluster samples and construct sample correlation network. The comparison with CSmiR shows that with using priori information of miRNA targets, Scan mostly has better or similar performance compared with CSmiR^16^ in inferring cell-specific miRNA regulation from the chronic myelogenous leukemia dataset. Furthermore, without using priori information of miRNA targets, the performance of Scan is mostly better than or comparable to the performance of CSmiR in inferring tissue-specific miRNA regulation from the breast cancer dataset.

Altogether, Scan provides a useful way to identify sample-specific miRNA regulation in human cancers, and can help to elucidate miRNA regulation mechanisms at the resolution of individual samples.

## Results

### <u>S</u>ample-spe<u>c</u>ific miRNA regul<u>a</u>tio<u>n</u> (Scan)

To investigate miRNA regulation at the resolution of single samples, we develop an approach called Scan to infer sample-specific miRNA regulation from bulk and single-cell RNA-sequencing data (**Fig. 1**). The bulk RNA-sequencing data of 690 breast cancer (BRCA) tissues is obtained from The Cancer Genome Atlas (TCGA)^26^ project, and the single-cell RNA-sequencing data of 19 half K562 cells (the first human chronic myeloid leukemia cell line) is from the Gene Expression Omnibus (GEO)^27^ database (Methods). The optional priori information of miRNA-mRNA interactions is from two well-known databases: TargetScan v8.0^28^ and ENCORI^29^ (Methods).

**Fig. 1.**
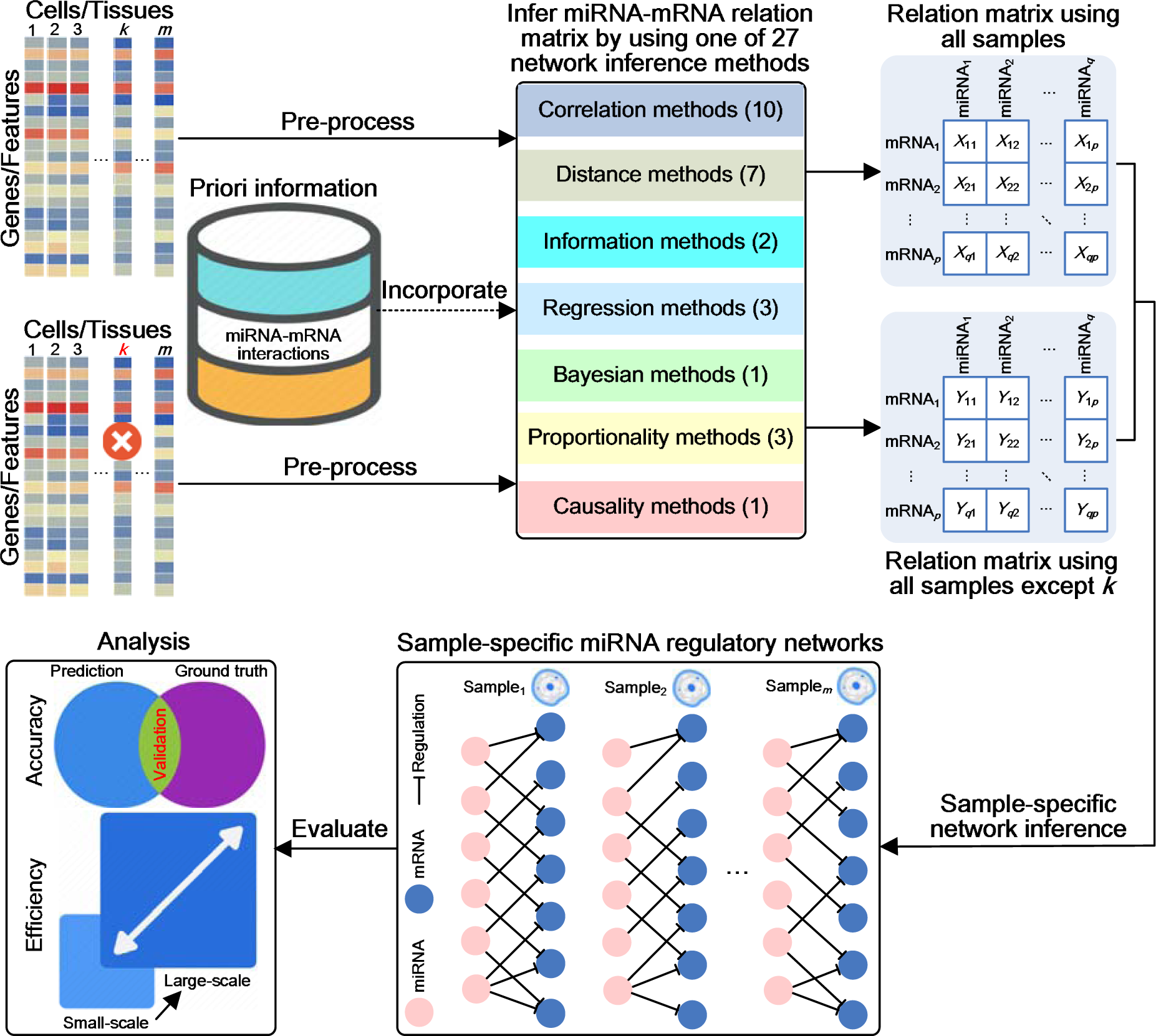
Schematic illustration of Scan. Given bulk or single-cell RNA-sequencing data with or without priori information of miRNA-mRNA interactions, Scan applies one of 27 network inference methods covering seven types (Correlation, Distance, Information, Regression, Bayesian, Proportionality and Causality) to construct miRNA-mRNA relation matrix. By using one of 27 network inference methods, Scan constructs two miRNA-mRNA relation matrices (one for all samples and the other for all samples except the *k*-th sample). Then for sample (cell or tissue) *k*, Scan conducts sample-specific network inference from the two constructed miRNA-mRNA relation matrices to infer a miRNA regulatory network specific to the sample *k*. In total, Scan can identify *m* sample-specific miRNA regulatory networks across *m* samples (one network for one sample).

Given bulk or single-cell RNA-sequencing data with or without priori information of miRNA-mRNA interactions, Scan apply one of 27 network inference methods (Methods) spanning seven types (including Correlation, Distance, Information, Regression, Bayesian, Proportionality and Causality) to infer miRNA-mRNA relation matrix. For each sample (cell or tissue), Scan needs to infer two miRNA-mRNA relation matrices (one for all samples and the other for all samples except the sample of interest). Based on the two identified miRNA-mRNA relation matrices, Scan further adapts one strategy (statistical perturbation or linear interpolation) to infer miRNA regulatory network specific to the sample of interest (Methods).

### Cell-specific miRNA regulation in K562

Using the single-cell miRNA-mRNA co-sequencing data in K562, we apply Scan to infer cell-specific miRNA regulation across 19 half K562 cells. Scan firstly uses 27 network inference methods to infer miRNA-mRNA relation matrices. For each cell, each network inference method infers two miRNA-mRNA relation matrices (one for all cells and the other for all cells except the cell of interest). Based on the identified miRNA-mRNA relation matrices, Scan further applies two strategies Scan.interp (using linear interpolation strategy) and Scan.perturb (using statistical perturbation strategy) to construct cell-specific miRNA regulatory networks across 19 half K562 cells. To systematically compare the performance of different combinations of network inference methods and strategies used, we have analyzed a total of 1026 networks: 27 (network inference methods) times 2 (strategies) times 19 (cells). We have found that using different network inference methods results in different number of cell-specific miRNA-mRNA interactions, and Dcor^30^ has the largest number of miRNA-mRNA interactions predicted (**Supplementary** Fig. 1a and 2a in **Supplementary File 1**). Furthermore, when priori information of miRNA targets is used, more cell-specific miRNA regulatory networks are found to have their node degree obey a power law distribution and thus are regarded as scale-free networks (**Supplementary** Fig. 1b and 2b in **Supplementary File 1**). Interestingly, with or without using priori information of miRNA targets, the node degree of the identified cell-specific miRNA regulatory networks by Dcor does not follow a power law distribution. In terms of the predicted miRNA-mRNA interactions, the similarity of the majority of cell pairs is less than 0.50 (**Supplementary** Fig. 1c and 2c in **Supplementary File 1**), indicating the heterogeneity of miRNA regulation for each K562 cell.

When Scan.interp is used for inferring cell-specific miRNA regulation, using Lasso^31^ for strength calculation gives the best final performance in terms of accuracy and using Bcor^32^ leads to the highest efficiency (**Fig. 2**). Overall, using Canberra^33^ has the largest overall rank score when using Scan.interp for inferring cell-specific miRNA regulation. When Scan.perturb is used for inferring cell-specific miRNA regulation, using Lasso also has the best final performance in terms of accuracy whereas using Bcor also leads to the highest efficiency (**Fig. 2**). Overall, using Chebyshev^34^ shows the best overall performance when using Scan.perturb for inferring cell-specific miRNA regulation. This result indicates that in terms of accuracy and efficiency, Lasso and Bcor are consistent in the K562 dataset regardless of which strategy is used for inferring cell-specific miRNA regulation. Although the best network inference method (in terms of overall rank score) is different between Scan.interp and Scan.perturb, the network inference methods which lead to the best final result are all distance based network inference methods. Furthermore, in terms of efficiency, the rank scores obtained by using 27 network inference methods between Scan.interp and Scan.perturb are consistent (*p*-value less than 2.22E-16 with Cohen’s kappa). However, in terms of accuracy and overall, the rank scores obtained by using 27 network inference methods between Scan.interp and Scan.perturb are not consistent (*p*-values are 0.53 and 0.69 with Cohen’s kappa, respectively). This result suggests that the performance of the network inference methods with the K562 dataset is closely related to the selected strategy for sample-specific miRNA regulatory networks.

**Fig. 2.**
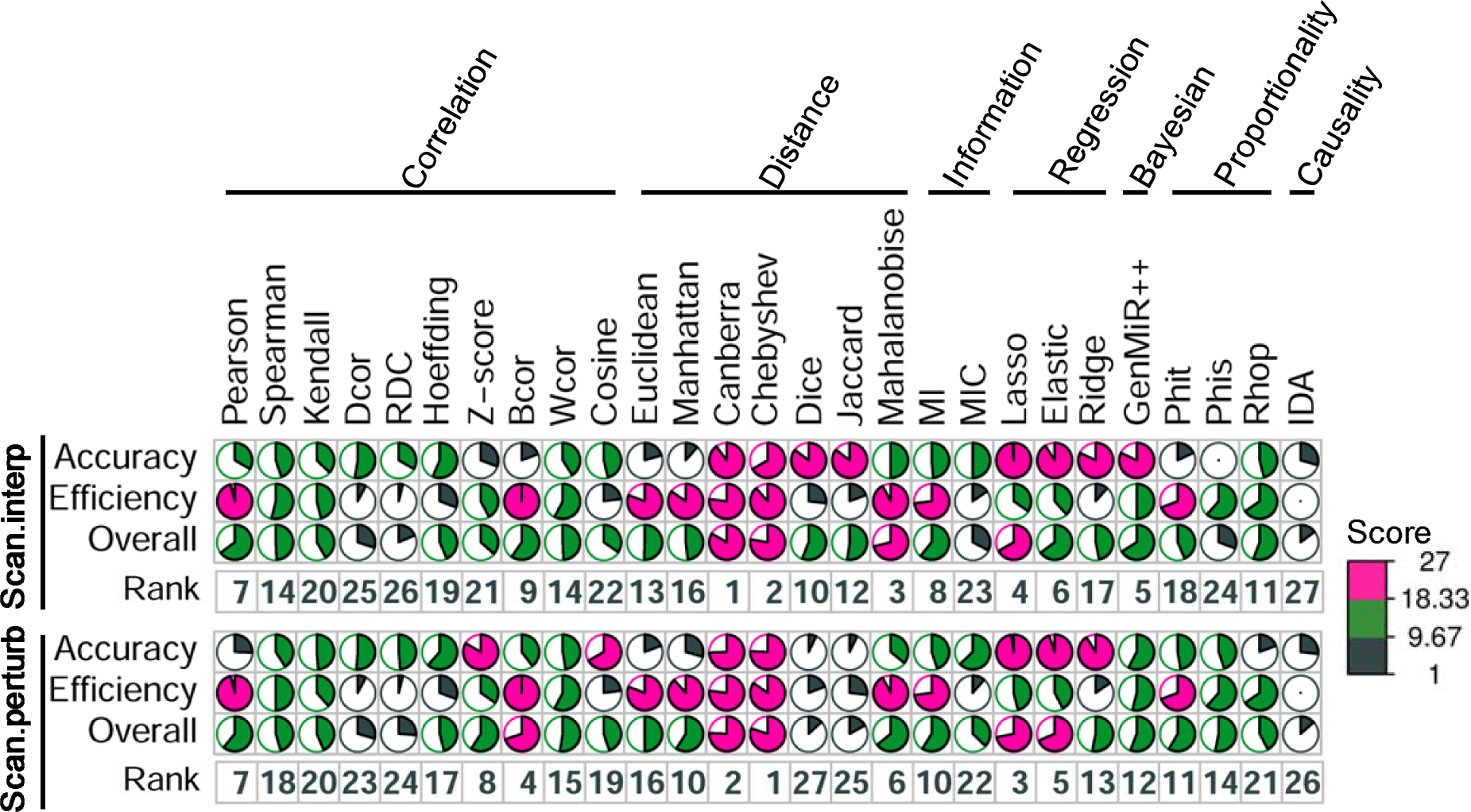
Performance of each network inference method in K562 using Scan.interp and Scan.perturb. Scores for each network inference method are calculated to evaluate the performance of predicting cell-specific miRNA regulation in terms of accuracy, efficiency and overall. Higher scores and smaller ranks denote better performance. The rank is visualized as a clock where clockwise direction indicates descending order.

### _r_Tissue-specific miRNA regulation in BRCA

Using the bulk miRNA and mRNA expression data in BRCA, we apply Scan to infer tissue-specific miRNA regulation across 690 BRCA tissues. Firstly, Scan uses 22 network inference methods (five methods are discarded due to long runtime or large memory usage) to infer miRNA-mRNA relation matrices. For each tissue, each network inference method infers two miRNA-mRNA relation matrices (one for all tissues and the other for all tissues except the tissue of interest). For Scan.interp and Scan.perturb, Scan generates tissue-specific miRNA regulatory networks across 690 BRCA tissues based on the identified miRNA-mRNA relation matrices. To systematically evaluate the performance of different combinations of network inference methods and strategies used, we have analyzed a total of 30,360 networks: 22 (network inference methods) times 2 (strategies) times 690 (tissues). We have discovered that using different network inference methods leads to different number of tissue-specific miRNA-mRNA interactions (**Supplementary** Fig. 3a and 4a in **Supplementary File 1**). Moreover, when priori information of miRNA targets is used, more tissue-specific miRNA regulatory networks are found to have their node degree follow a power law distribution and thus are regarded as scale-free networks (**Supplementary** Fig. 3b and 4b in **Supplementary File 1**). In terms of the predicted miRNA-mRNA interactions, the similarity of most of tissue pairs is also less than 0.50 (**Supplementary** Fig. 3c and 4c in **Supplementary File 1**), revealing the heterogeneity of miRNA regulation for each BRCA tissue.

When Scan.interp is used for inferring tissue-specific miRNA regulation, using GenMiR++^35^ for strength calculation gives the best final results in terms of accuracy and whereas using Bcor^32^ has the highest efficiency (**Fig. 3**). Overall, using Phis^36^ has the largest overall rank score when using Scan.interp for inferring tissue-specific miRNA regulation. When Scan.perturb is used for inferring tissue-specific miRNA regulation, using Rhop^36^ gives the best final results in terms of accuracy whereas using Bcor also has the highest efficiency (**Fig. 3**). Overall, using Rhop^36^ displays the best performance when using Scan.perturb for inferring tissue-specific miRNA regulation. This result suggests that in terms of efficiency, Bcor is consistent in the BRCA dataset regardless of which strategy is used for inferring tissue-specific miRNA regulation. Although the best network inference method (in terms of overall rank score) is different between Scan.interp and Scan.perturb, the network inference methods which lead to the best final result are all proportionality based network inference methods. Moreover, in terms of efficiency, the rank scores obtained by using 22 network inference methods between Scan.interp and Scan.perturb are consistent (*p*-value less than 2.22E-16 with Cohen’s kappa). However, in terms of accuracy and overall, the rank scores obtained by using 22 network inference methods between Scan.interp and Scan.perturb are not consistent (*p*-values are 0.58 and 0.75 with Cohen’s kappa, respectively). This result implies that the performance of the network inference methods with the BRCA dataset is also closely associated with the selected strategy.

**Fig. 3.**
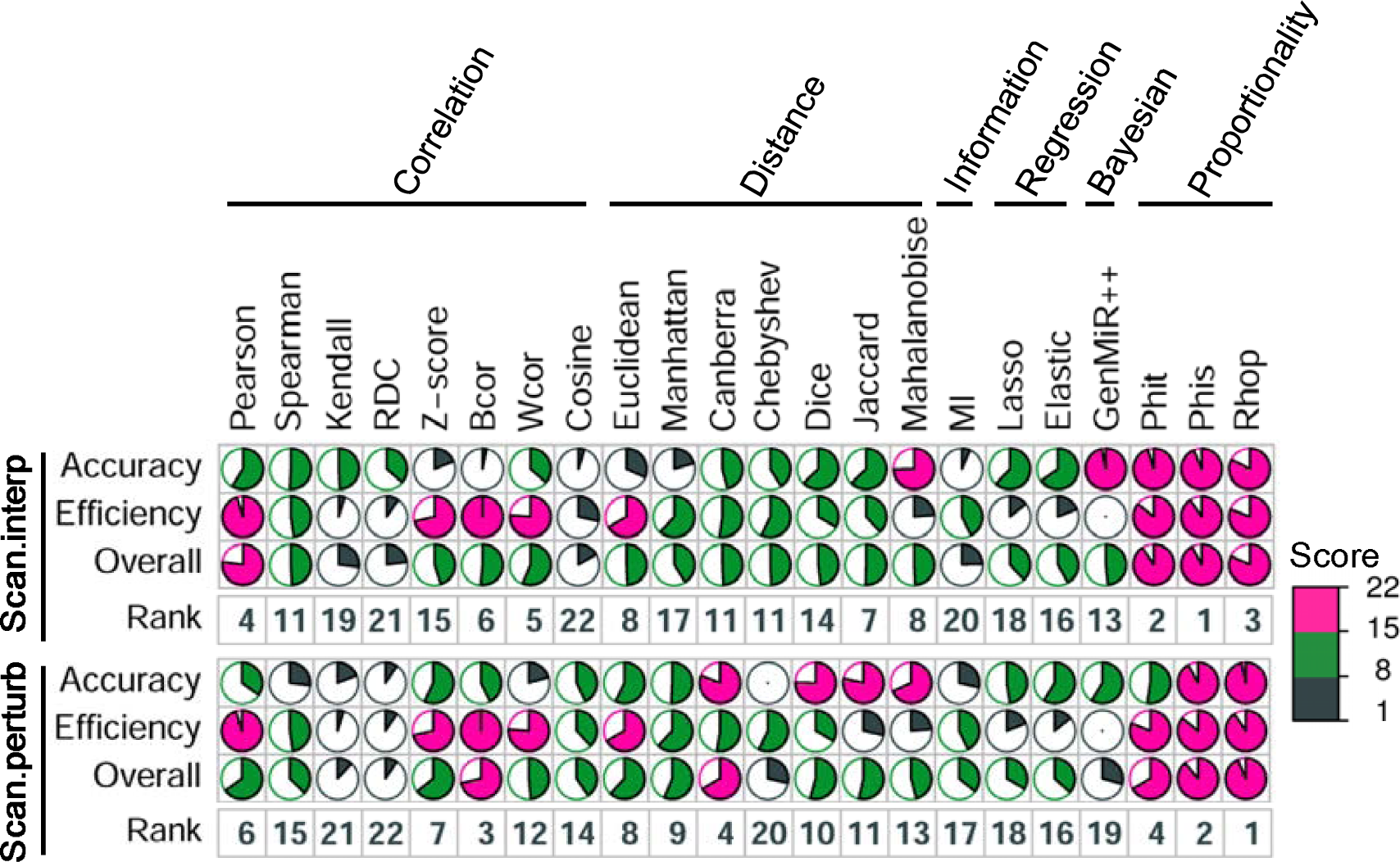
Performance of each network inference method in BRCA using Scan.interp and Scan.perturb. Scores for each network inference method are calculated to evaluate the performance of predicting tissue-specific miRNA regulation in terms of accuracy, efficiency and overall. Higher scores and smaller ranks represent better performance. The rank is visualized as a clock where clockwise direction indicates descending order.

### Priori information can improve the accuracy of miRNA target prediction

To understand whether priori information of miRNA targets can improve the accuracy of miRNA target prediction, we compare the average percentage of validated miRNA targets predicted by Scan with and without using priori information of miRNA targets. We find that incorporating priori information of miRNA targets generally results in a larger average percentage of validated miRNA targets in both K562 and BRCA datasets (**Fig. 4**). Moreover, the higher the confidence of the priori information is, generally the larger the average percentage of validated miRNA targets in both K562 and BRCA datasets is (**Fig. 4**). This finding demonstrates that the priori information of miRNA targets can improve the accuracy of miRNA target prediction.

**Fig. 4.**
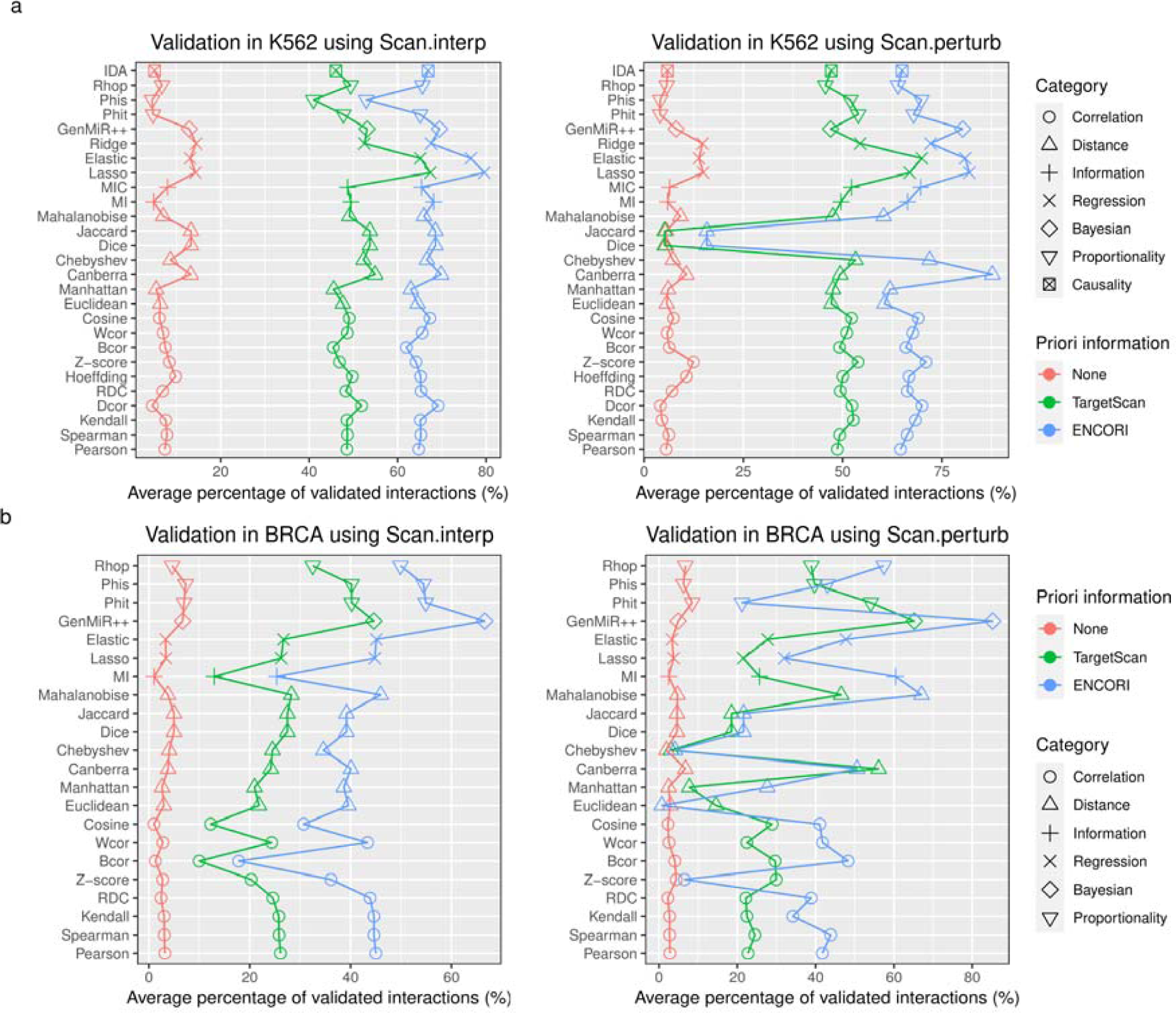
Validation of sample-specific miRNA regulations with and without using priori information. **a** Validation in the K562 dataset using Scan.interp and Scan.perturb. **b** Validation in the BRCA dataset using Scan.interp and Scan.perturb.

### Clusters of K562 cells and BRCA tissues

With the K562 dataset, when Scan.interp and Scan.perturb are used together with Canberra and Chebyshev respectively, the best result in terms of overall rank score is achieved. For the BRCA dataset, when Scan.interp and Scan.perturb are used together with Phis and Rhop respectively, the best result in terms of overall rank score is obtained. As a result, we select the optimal combinations, Scan.interp_Canberra (Scan.interp with Canberra) and Scan.perturb_Chebyshev (Scan.perturb with Chebyshev) for the K562 dataset to infer cell-specific miRNA regulatory networks, and Scan.interp_Phis (Scan.interp with Phis) and Scan.perturb_Rhop (Scan.perturb with Rhop) for the BRCA dataset to identify tissue-specific miRNA regulatory networks. We use the identified cell-specific (or tissue-specific) miRNA regulatory networks without using priori information to construct cell-cell (or tissue-tissue) similarity matrices. Based on the constructed cell-cell and tissue-tissue similarity matrices, we further conduct hierarchical clustering analysis to cluster K562 cells and BRCA tissues, respectively (Methods).

With the K562 dataset, we have three distinct clusters of K562 cells (**Fig. 5a and 5b**). With the BRCA dataset, we obtain five clear clusters of BRCA tissues (**Supplementary** Fig. 5a and 5b in **Supplementary File 1, Supplementary Data 1**). It is noted that the different clustering results of K562 cells or BRCA tissues are explained by using different network inference methods and different strategies. Since gene regulatory networks (e.g. miRNA regulatory networks) are regarded as stable forms to characterize a cell or a tissue^37,38^, our clustering results based on the identified cell-specific or tissue-specific miRNA regulatory networks may help to identify novel cell or tissue subtypes.

**Fig. 5.**
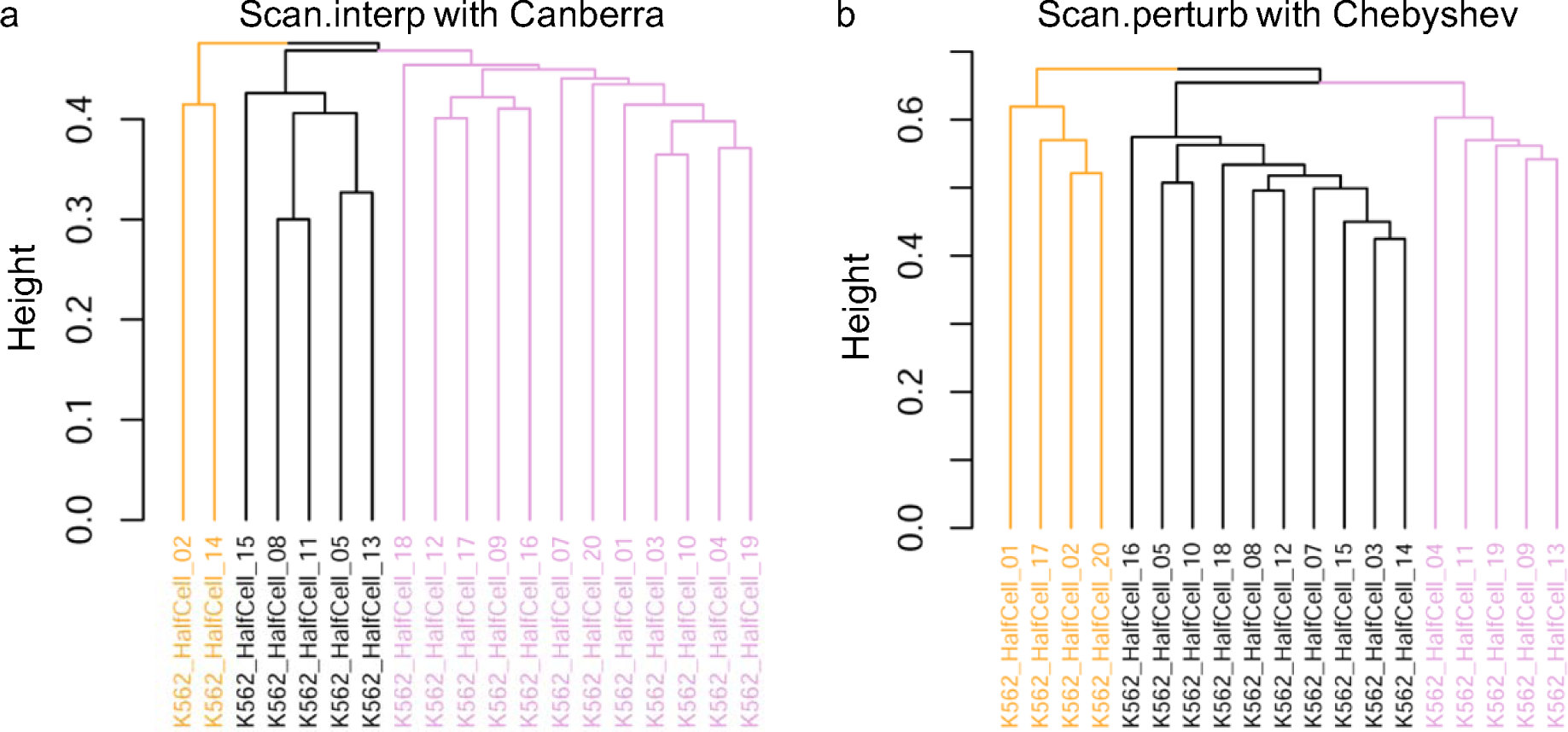
Clusters of K562 cells. **a** Clusters of K562 cells, and Scan.interp with Canberra is used to identify cell-specific miRNA regulatory networks. **b** Clusters of K562 cells, and Scan.perturb with Chebyshev is used to infer cell-specific miRNA regulatory networks. Each color denotes a cluster.

### Correlation networks of K562 cells and BRCA tissues

Similar to the clustering of K562 cells and BRCA tissues, we also select the four optimal combinations (Scan.interp_Canberra, Scan.perturb_Chebyshev, Scan.interp_Phis, Scan.perturb_Rhop) without using priori information to infer cell-specific and tissue-specific miRNA regulatory networks respectively for the two datasets. We use the inferred cell-specific (or tissue-specific) miRNA regulatory networks to generate cell-cell (or tissue-tissue) similarity matrices. Based on the generated cell-cell and tissue-tissue similarity matrices, we further construct correlation networks of K562 cells and BRCA tissues respectively (Methods).

We have constructed two correlation networks for the K562 cells and two correlation networks for the BRCA tissues (**Supplementary Data 2**). Based on the identified cell-specific miRNA regulatory networks by Scan.interp_Canberra and Scan.perturb_Chebyshev, the numbers of cell-cell correlation pairs are 171 and 22 respectively for the K562 dataset, Based on the identified tissue-specific miRNA regulatory networks by Scan.interp_Phis and Scan.perturb_Rhop, the numbers of tissue-tissue correlation pairs are 51,769 and 8805 respectively for BRCA dataset. Network topological analysis reveals that the node degree of the identified cell-cell correlation network (10 nodes and 22 edges) by Scan.perturb_Chebyshev obeys a power law distribution (*p*-value more than 0.99 with Kolmogorov-Smirnov test), and the node degrees of two identified tissue-tissue correlation networks by Scan.interp_Phis and Scan.perturb_Rhop also follow power law distributions (*p*-values more than 0.85 with Kolmogorov-Smirnov test). Additionally, the identified cell-cell correlation network (19 nodes and 171 edges) by Scan.interp_Canberra is a fully connected network. This result indicates that the identified correlation networks of cells or tissues tend to be scale-free or fully connected networks.

### Dynamic and conservative miRNA regulation across K562 cells and BRCA tissues

We also choose the four optimal combinations (Scan.interp_Canberra, Scan.perturb_Chebyshev, Scan.interp_Phis, Scan.perturb_Rhop) without using priori information to infer miRNA regulatory networks specific to K562 cells and BRCA tissues. Based on the identified cell-specific or tissue-specific miRNA regulatory networks, we further investigate the dynamic and conservative miRNA regulation across K562 cells and BRCA tissues (Methods).

With both K562 and BRCA dataset, the numbers of dynamic and conservative miRNA-mRNA interactions identified tend to be varying (**Supplementary Data 3**). With the K562 dataset, the number of dynamic miRNA-mRNA interactions identified by Scan.interp_Canberra (13786) is smaller than that identified by Scan.perturb_Chebyshev (77627), but the number of conservative miRNA-mRNA interactions identified by Scan.interp_Canberra (6490) is larger than that identified by Scan.perturb_Chebyshev (3213). With the BRCA dataset, the numbers of dynamic and conservative miRNA-mRNA interactions identified by Scan.interp_Phis (82850 and 29053, respectively) are larger than those identified by Scan.perturb_Rhop (5549 and 36, respectively). Moreover, for Scan.interp_Canberra, Scan.perturb_Chebyshev, Scan.interp_Phis and Scan.perturb_Rhop, the number of miRNAs involved in the dynamic miRNA regulation (45, 212, 163 and 98, respectively) is larger than or equal to that of the conservative miRNA regulation (8, 212, 37 and 25, respectively).

With the K562 dataset, the node degrees of the identified dynamic and conservative miRNA regulatory networks by Scan.interp_Canberra do not follow power law distributions (*p*-values are 2.41E-05 and 3.77E-04 with Kolmogorov-Smirnov test, respectively), but the node degrees of the identified dynamic and conservative miRNA regulatory networks by Scan.perturb_Chebyshev obey power law distributions (*p*-values are 1.00 and 0.99 with Kolmogorov-Smirnov test, respectively). With the BRCA dataset, the node degrees of the identified dynamic miRNA regulatory networks by Scan.interp_Phis and the identified conservative miRNA regulatory networks by Scan.perturb_Rhop follow power law distributions (*p*-values are 0.21 and 1.00 with Kolmogorov-Smirnov test, respectively). However, with the BRCA dataset, the node degrees of the identified conservative miRNA regulatory networks by Scan.interp_Phis and the identified dynamic miRNA regulatory networks by Scan.perturb_Rhop do not obey power law distributions (*p*-values are 2.77E-33 and 9.47E-05 with Kolmogorov-Smirnov test, respectively).

In addition, the numbers of dynamic and conservative hub miRNAs by Scan.interp and Scan.perturb are also different (**Supplementary Data 4**). With the K562 dataset, the number of dynamic and conservative hub miRNAs identified by Scan.interp_Canberra (16 and 1, respectively) is larger than those identified by Scan.perturb_Chebyshev (0 and 0, respectively). With the BRCA dataset, the numbers of dynamic and conservative hub miRNAs identified by Scan.interp_Phis (44 and 15, respectively) are larger than those identified by Scan.perturb_Rhop (39 and 0, respectively). Furthermore, with the K562 dataset, 4 out of 16 dynamic hub miRNAs and 1 out of 1 conservative hub miRNA identified by Scan.interp_Canberra are confirmed to be CML-related miRNAs. With the BRCA dataset, 38 out of 44 dynamic hub miRNAs and 15 out of 15 conservative hub miRNAs identified by Scan.interp_Phis are confirmed to be BRCA-related miRNAs, and 39 out of 39 dynamic hub miRNAs identified by Scan.perturb_Rhop are validated to be BRCA-related miRNAs.

Altogether, the above results suggest that the findings of the dynamic and conservative miRNA regulation across K562 cells and BRCA tissues depend on the selected network inference methods and strategies.

### Performance comparison

To evaluate the effectiveness of Scan, we compare the performance of Scan (Scan.interp and Scan.perturb) with CSmiR^16^ on the K562 and BRCA datasets. Here, CSmiR is the first method to explore miRNA regulation at a single-cell resolution level, so we use it as the baseline for comparison. With the K562 dataset, without using priori information of miRNA targets, Scan.interp with 8 network inference methods and Scan.perturb with 7 network inference methods perform better than CSmiR in terms of accuracy (**Fig. 6a**). After integrating priori information of miRNA targets from TargetScan, Scan.interp with 14 network inference methods and Scan.perturb with 18 network inference methods perform better than CSmiR in terms of accuracy, and after integrating priori information of miRNA targets from ENCORL, Scan.interp with 21 network inference methods and Scan.perturb with 20 network inference methods perform better than CSmiR in terms of accuracy (**Fig. 6a**). This result shows that in terms of accuracy, Scan mostly performs better than or comparable to CSmiR with the K562 dataset when using priori information of miRNA targets. Furthermore, in terms of efficiency, both Scan.interp and Scan.perturb with all of the network inference methods perform better than CSmiR (**Fig. 6b**).

**Fig. 6.**
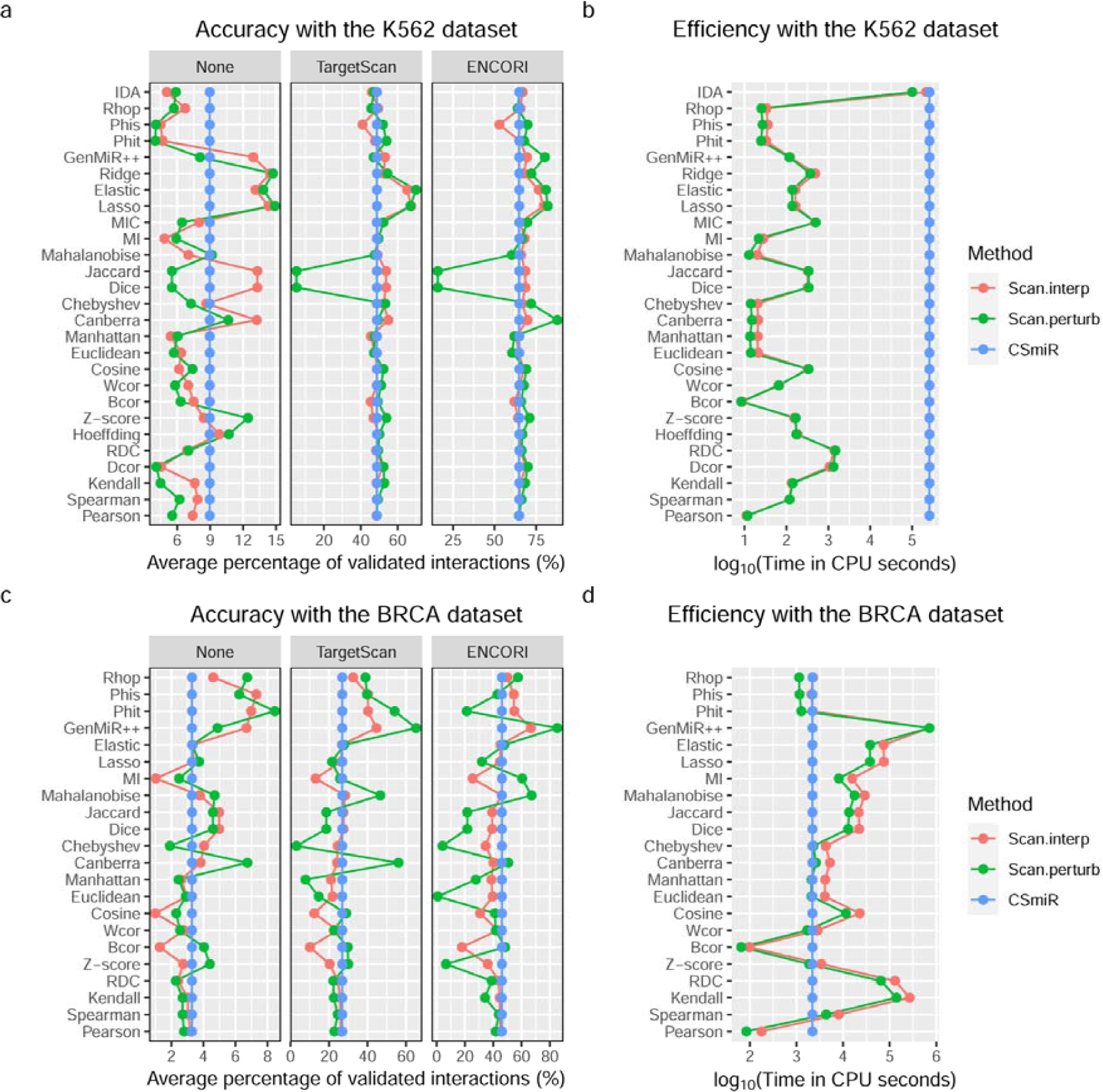
Performance comparison in inferring sample-specific miRNA regulation between Scan.interp, Scan.perturb and CSmiR. **a** Accuracy comparison with the K562 dataset. **b** Efficiency comparison with the K562 dataset. **c** Accuracy comparison with the BRCA dataset. **d** Efficiency comparison with the BRCA dataset.

With the BRCA dataset, without using priori information of miRNA targets, Scan.interp with 11 network inference methods and Scan.perturb with 12 network inference methods perform better than CSmiR in terms of accuracy (**Fig. 6c**). After integrating priori information of miRNA targets from TargetScan, Scan.interp with 7 network inference methods and Scan.perturb with 10 network inference methods perform better than CSmiR in terms of accuracy, and after integrating priori information of miRNA targets from ENCORL, Scan.interp with 4 network inference methods and Scan.perturb with 7 network inference methods perform better than CSmiR in terms of accuracy (**Fig. 6c**). This result shows that in terms of accuracy, Scan mostly performs better than or comparable to CSmiR with the BRCA dataset when without using priori information of miRNA targets. In addition, in terms of efficiency, Scan.interp with 3 network inference methods and Scan.perturb with 9 network inference methods perform better than CSmiR (**Fig. 6d**).

The above performance comparison suggests that in the small-scale K562 dataset, Scan with priori information generally performs better than or comparable to CSmiR in terms of accuracy and efficiency. In the large-scale BRCA dataset, Scan without using priori information generally performs better than or comparable to CSmiR in terms of accuracy, but in terms of efficiency, only a few instances of Scan (Scan with a few network inference methods) perform better than CSmiR. The inconsistent result of performance comparison with the two datasets implies that the performance of Scan is more likely to be data-specific, and it is necessary to select optimal network inference methods for new data.

## Discussion

It is well-established that miRNAs are important regulators of gene expression, and their dysregulations can lead to the occurrence and development of complex human diseases, including cancers. Yet, the research on miRNA regulation at the resolution of single samples (cells or tissues) is still limited. Here, we present the Scan framework, and further show the effectiveness of it in inferring sample-specific miRNA regulation. By applying Scan into bulk and single-cell RNA-sequencing data, we have discovered that adding priori information of miRNA targets can improve the accuracy of miRNA target prediction. By instantiating the Scan framework with 27 network inference methods, we have found that the performance of Scan instantiated with different network inference methods exhibits to be data-specific. In addition, the identified sample-specific miRNA regulatory network by Scan can be used for downstream analysis, e.g. clustering samples and constructing sample correlation network. The freely available framework Scan provides a useful method for exploring miRNA regulation at the resolution of single samples.

For constructing miRNA-mRNA relation matrix, Scan includes 27 network inference methods spanning seven types (Correlation, Distance, Information, Regression, Bayesian, Proportionality and Causality). It is noted that other types of computational methods (e.g. deep learning^39^, probabilistic modeling^40^) can be also plugged into Scan for generating miRNA-mRNA relation matrix. To improve the generalization ability of Scan, it is our plan to add more types of network inference methods in future.

When applying Scan to a new data, user can choose one or multiple network inference methods according to the rank score in terms of accuracy or efficiency or overall. For example, for the K562 dataset, we can select network inference method Canberra to be used with Scan.interp and Chebyshev with Scan.perturb in terms of overall rank score. But for the BRCA dataset, we can select network inference method Phis to be used with Scan.interp and Rhop with Scan.perturb in terms of overall rank score.

We provide two strategies, Scan.interp (linear interpolation strategy) and Scan.perturb (statistical perturbation strategy) to infer miRNA regulation specific to single cells or tissues. Generally, using the same network inference method, Scan.interp can generate larger cell-specific or tissue-specific miRNA regulatory networks than those by Scan.perturb. This can be explained by the fact that Scan.interp infers cell-specific miRNA regulatory networks with differential values (e.g. differential correlation, distance or regression) between all samples and all samples except sample *k*, but Scan.perturb identifies differential miRNA regulatory networks between the networks of using all samples and the networks of using all samples except sample *k* (see details in Methods). Moreover, the runtime of Scan.interp and Scan.perturb with the K562 and BRCA datasets is similar. For new data, we suggest use both Scan.interp and Scan.perturb for comprehensively exploring miRNA regulation specific to samples.

To infer sample-specific miRNA regulation, we have applied Scan into bulk and single-cell RNA-sequencing data across tumor cells or tissues. Certainly, Scan is also applicable in bulk and single-cell RNA-sequencing data across healthy cells or tissues. In future, it will be a meaningful direction to reveal dynamic and conservative miRNA regulation between matched tumor and healthy cells or tissues. Moreover, with the advancement of spatial RNA-sequencing technology^41^, Scan is also a potential method to explore the heterogeneity of miRNA regulation between different spatial positions from spatial RNA-sequencing data.

When Scan is applied to large-scale transcriptomics data, the process of inferring miRNA-mRNA relation matrix using network inference methods will be computationally intensive. This is a common issue of existing computational methods, including Scan. To alleviate the issue, users can allocate more CPU cores to identify miRNA-mRNA relation matrix in parallel. Furthermore, users can also use priori information of samples to divide samples into several subtypes, and then focus on investigating sample-specific miRNA regulation across cells or tissues of each subtype. Finally, selecting a fast network inference method (e.g. Pearson and Bcor) is also a feasible way to quickly infer sample-specific miRNA regulation from large-scale transcriptomics data.

In addition to studying miRNA regulation at single-sample resolved level, Scan also can be used for other types of gene regulation specific to individual samples, e.g. transcriptional regulation, long non-coding RNA regulation, circular RNA regulation and PIWI-interacting RNA regulation.

Since miRNAs play important roles in tumor microenvironments, it is important for us to understand the regulation of miRNAs in tumor cells or tissues. In-depth investigation of miRNA regulation specific to individual tumor cells or tissues will help to figure out the heterogeneity of tumor microenvironments and discover novel subtypes of tumor cells or tissues.

## Methods

### Bulk and single-cell RNA-sequencing data

From TCGA^26^, we have obtained the expression data of 894 miRNAs and 19,068 mRNAs in 690 matching breast cancer tissues. Since epithelial-mesenchymal transition (EMT) is closely related to the development, progression and metastasis of breast cancer^42–45^, we obtain a list of 315 EMT signatures^46^ (**Supplementary Data 5**) and further divide the matched 690 breast cancer tissues into four EMT types (epithelial, intermediate epithelial, intermediate mesenchymal and mesenchymal) by using GSVA R package^47^. As a result, the numbers of breast cancer tissues belonging to epithelial, intermediate epithelial, intermediate mesenchymal and mesenchymal types are 491, 107, 46 and 46, respectively (**Supplementary Data 6**). By using the *limma-trend* approach in limma R package^48^, we have identified 163 miRNAs and 5801 mRNAs that are differentially expressed between epithelial and mesenchymal type (adjusted *p*-value < 0.01, fold change > 1.5) (**Supplementary Data 7**). Here, the *p*-values are adjusted by the Benjamini–Hochberg (BH) method^49^. Therefore, the input bulk RNA-sequencing data used in our case study includes the expression data of 163 miRNAs and 5801 mRNAs in 690 breast cancer tissues.

The K562 single-cell RNA-sequencing data (the accession number is GSE114071 in GEO database^27^) includes the expression data of 2822 miRNAs and 21,704 mRNAs in 19 half K562 cells (the first human chronic myeloid leukemia cell line). In the single-cell RNA-sequencing data, we calculate the average expression values of the duplicate miRNAs or mRNAs as their ultimate expression values. As a feature selection step, we discard the miRNAs or mRNAs with constant expression values, and we are interested in the high expression miRNAs or mRNAs with mean values greater than the median of mean values of all non-constant expression miRNAs or mRNAs. Moreover, the reserved miRNA data and mRNA expression data are then log-transformed. As a result, the input single-cell RNA-sequencing data used in our case study includes the expression data of 212 miRNAs and 7680 mRNAs in 19 half K562 cells.

### Priori information of miRNA targets

To improve the prediction of sample-specific miRNA regulation, the miRNA target information in TargetScan v8.0^28^ and ENCORI^29^ (the pilot version is starBase) is used as priori information for Scan. From TargetScan, a list of 235,109 predicted miRNA-mRNA interactions has been obtained. To identify miRNA-mRNA interactions, ENCORI provides seven prediction tools (PITA^50^, RNA22^51^, miRmap^52^, DIANA-microT^53^, miRanda^54^, PicTar^55^ and TargetScan^28^) for users. To obtain high-confidence miRNA-mRNA interactions, we only retain the miRNA-mRNA interactions predicted by at least five prediction algorithms. In total, a list of 55,343 high-confidence miRNA-mRNA interactions are obtained from ENCORI.

### Network inference methods

We select 27 network inference methods (**Supplementary Data 8**) for constructing miRNA-mRNA relation matrix: Pearson^56^, Spearman^57^, Kendall^58^, Distance correlation (Dcor)^30^, Random Dependence Coefficient (RDC)^59^, Hoeffding’s D statistics (Hoeffding)^60^, Z-score^61^, Biweight midcorrelation (Bcor)^32^, Weighted rank correlation (Wcor)^62^, Cosine^63^, Euclidean^64^, Manhattan^65^, Canberra^33^, Chebyshev^34^, Dice^66^, Jaccard^67^, Mahalanobise^68^, Mutual Information (MI)^69^, Maximal Information Coefficient (MIC)^70^, Lasso^31^, Elastic^31^, Ridge^31^, GenMiR++^35^, *φ* (Phit)^36^, *φ*_s_ (Phis)^36^, *ρ*_p_ (Rhop)^36^, and Intervention calculus when the Directed acyclic graph is Absent (IDA)^71^. These methods can be divided into seven types: Correlation,

Distance, Information, Regression, Bayesian, Proportionality and Causality. Ten methods (Pearson, Spearman, Kendall, Dcor, RDC, Hoeffding, Z-score, Bcor, Wcor and Cosine) belong to the Correlation type, seven methods (Euclidean, Manhattan, Canberra, Chebyshev, Dice, Jaccard and Mahalanobise) belong to the Distance type, two methods (MI and MIC) are of the Information type, three methods (Lasso, Elastic and Ridge) belong to the Regression type, three methods (Phit, Phis and Rhop) belong to the Proportionality type, and GenMiR++ and IDA are of the Bayesian and Causality types, respectively.

### Sample-specific network inference

Given bulk or single-cell RNA-sequencing data with *m* samples, with or without priori information of miRNA-mRNA interactions, Scan constructs two miRNA-mRNA relation matrices (one for all samples and the other for all samples except the *k*-th sample of interest, *k* ∈[1, *m*]) by using one of the network inference methods. The two constructed miRNA-mRNA relation matrices (of *p* miRNAs and *q* mRNAs) for all samples and all samples except the *k*-th sample (denoted as *X*^(*k*)^ and *Y* ^(*k*)^ respectively) are as follows.

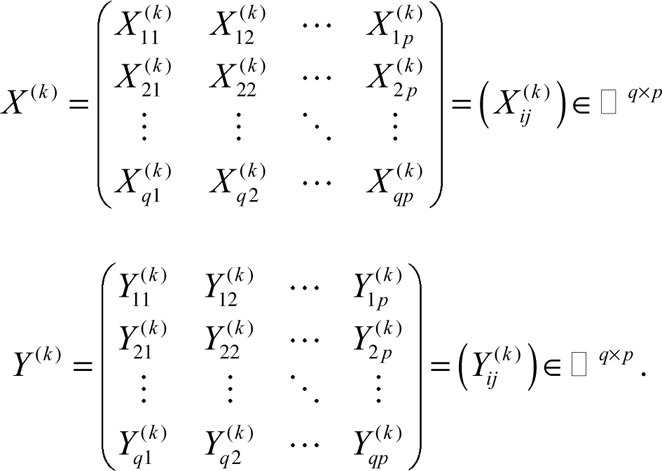

Where 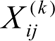 and 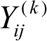 represent the connection between mRNA *i* and miRNA *j*.

For the network inference methods which are of the Correlation, Information, Regression, Bayesian, Proportionality and Causality types, larger absolute values of 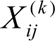 and 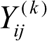 indicate higher connections between mRNA *i* and miRNA *j*. In contrast, for Distance based network inference methods, larger absolute values of 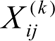 and 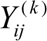 indicate weaker connections between mRNA *i* and miRNA *j*. To keep consistency between the constructed miRNA-mRNA relation matrices by different types of network inference methods, we transform the two miRNA-mRNA relation matrices (*X* ^(*k*)^ and *Y* ^(*k*^)) generated by Distance based methods into *X*^’(k)^ and *Y* ^’(*k*)^ as follows:

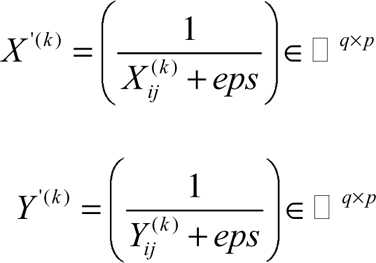

where *eps* refers to the precision of floating numbers (the value is 2.22E-16 in default).

After transformation, a larger absolute value in a relation matrix generated by any type of the network inference methods indicates a higher connection between a miRNA and a mRNA.

#### Linear interpolation strategy

Following the use of a network inference method of the Correlation, Information, Regression, Bayesian, Proportionality or Causality type to obtain the two constructed miRNA-mRNA relation matrices (*X* ^(*k*)^ and *Y* ^(*k*)^), Scan applies a linear interpolation strategy^22^ to estimate the miRNA-mRNA relation matrix *Z* ^(*k*)^ specific to sample *k* as follows:

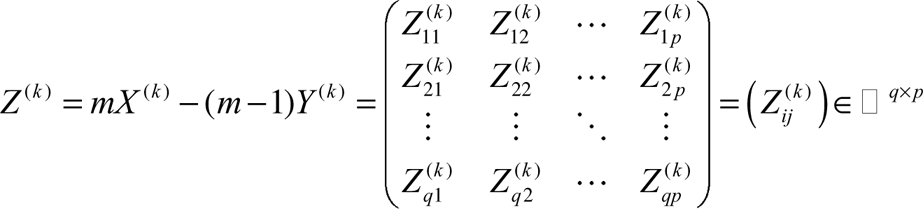

where *m* denotes the number of samples in bulk or single-cell RNA-sequencing data.

When a Distance based network inference method is used to obtain the two constructed miRNA-mRNA relation matrices (*X* ^’(*k*)^ and *Y* ^’(*k*)^), the miRNA-mRNA relation matrix *Z* ^(*k*)^ specific to sample *k* is estimated as:

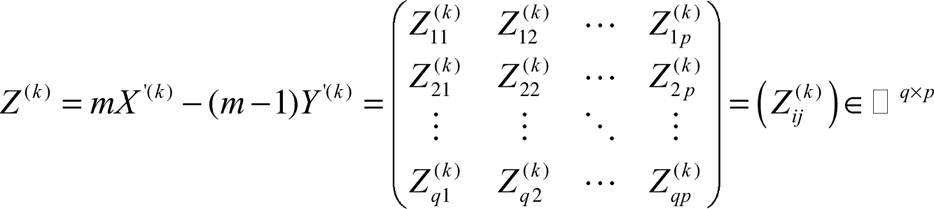

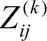 is further normalized as:

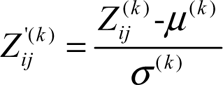

Where *μ* ^(*k*)^ and *_σ_* ^(*k*)^ denote the mean value and standard deviation of *Z* ^(*k*)^, and the normalized miRNA-mRNA relation matrix *Z* ^’(*k*)^ is:

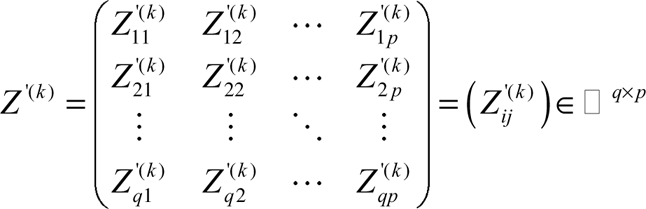

Each value of *Z* ^’(*k*)^ corresponds to a significance *p*-value. The *p*-value is calculated as follows:

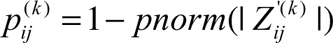

where | 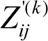 | is the absolute value of 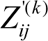, and the *pnorm* function is used to calculate the probability a random value from the standard normal distribution being less than | 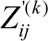 |.

Smaller value of 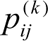 indicates that miRNA *j* is more likely to interact with mRNA *i* in sample *k*. If the value of 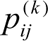 is less than a cutoff (e.g. 0.05), miRNA *j* is considered to interact with mRNA *i* in sample *k*. *Statistical perturbation strategy* For the Correlation, Distance, Information, Regression, Bayesian, Proportionality and Causality methods, each value of *X* ^(*k*)^ (*X* ^’(*k*)^) and *Y* ^(*k*)^ (*Y* ^’(*k*)^) corresponds to a significance *p*-value. The corresponding *p*-value matrices of *X* ^(*k*)^ (*X* ^’(*k*)^) and *Y* ^(*k*)^ (*Y* ^’(*k*)^) are denoted 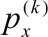 as and 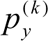, respectively.

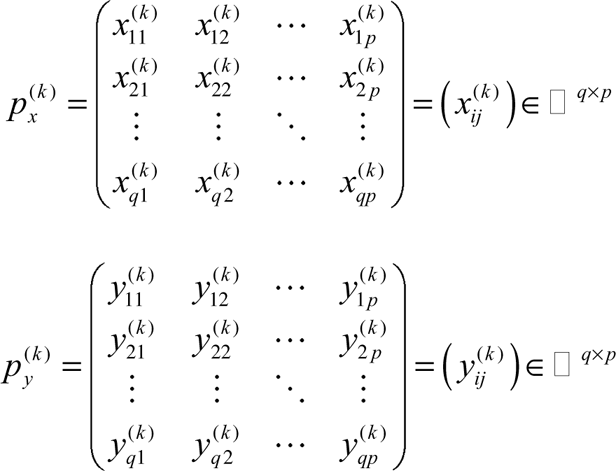

Smaller value of 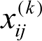 and 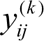 indicates that miRNA *j* is more likely to interact with mRNA *i* in all samples and all samples except *k*, respectively. Given a *p*-value cutoff (e.g. 0.05), we can obtain two zero-one matrices 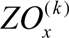 and 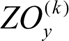 in all samples and all samples except *k*. Scan further uses a statistical perturbation strategy^21^ to calculate the miRNA-mRNA zero-one matrix *ZO*^(*k*)^ for sample *k* as follows:

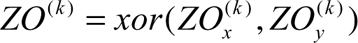

where *xor* is the XOR logical function. One value of 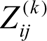 indicates that miRNA *j* interacts with mRNA *i* in sample *k*, and zero value of 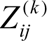 represents that miRNA *j* doesn’t interact with mRNA *i* in sample *k*.

In this work, the cutoff of significant *p*-value for both linear interpolation and statistical perturbation strategies is set to be 0.05 in default. By applying linear interpolation or statistical perturbation strategy, Scan can identify *m* sample-specific miRNA regulatory networks across *m* samples. Each sample-specific miRNA regulatory network is a directed graph where the direction of an edge is from a miRNA to a mRNA.

### Degree distribution analysis

The degree of a node in a sample-specific miRNA regulatory network is the number of connections with other nodes, and the degree distribution denotes the probability distribution of node degrees over the sample-specific miRNA regulatory network. If the node degree of a sample-specific miRNA regulatory network obeys a power law distribution, the network is considered as a scale-free network. In this work, we use the R package igraph^72^ to calculate the degree distribution of the identified sample-specific miRNA regulatory networks. The Kolmogorov-Smirnov (KS) test^73^ is used to determine whether the node degree of a sample-specific miRNA regulatory network obeys a power law distribution. If the *p*-value of the KS test is smaller than a cutoff (e.g. 0.05), the node degree of a sample-specific miRNA regulatory network does not obey a power law distribution, suggesting that the sample-specific miRNA regulatory network is not a scale-free network, and vice versa.

### Clustering analysis

We use sample-specific miRNA regulatory networks to compute the similarities between samples. In the context of sample-specific miRNA regulatory networks, the similarity between samples *a* and *b* is calculated in the following.

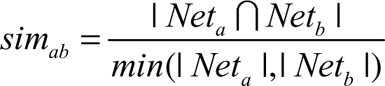

where *Net_a_* and *Net_b_* represent the miRNA regulatory networks specific to samples *a* and *b*, respectively, | *Net_a_* ∩ *Net_b_* | is the number of common miRNA-mRNA interactions between *Net_a_* and *Net_b_*, and *min*(| *Net_a_* |,| *Net_b_* |) denotes the smaller number of miRNA-mRNA interactions between *Net_a_* and *Net_b_*.

For *m* samples, the sample-sample similarity matrix *SM* (a symmetric matrix) is:

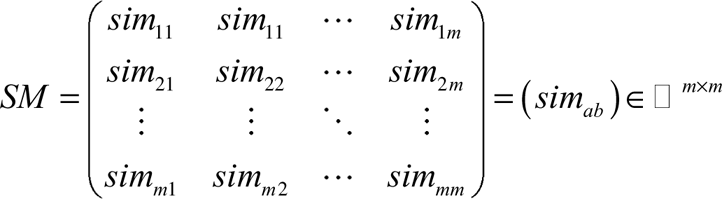

Based on the sample-sample similarity matrix *SM*, we further conduct clustering analysis of samples, e.g. hierarchical clustering analysis.

### Sample correlation network construction

In addition to the clustering analysis of samples, the sample-sample similarity matrix is also used for sample correlation network construction^74^. For each sample pair, a higher similarity value indicates that the pair of samples is more correlated with each other. We use an empirical similarity cutoff (e.g. 0.50) to infer whether two samples are correlated or not. In other words, if the similarity value between samples *a* and *b* is larger than the cutoff, samples *a* and *b* are correlated with each other. After assembling the correlated sample pairs, we can construct a sample correlation network.

### Dynamic and conservative analysis

The miRNA-mRNA interaction existing in only one cell or tissue are defined as a dynamic miRNA-mRNA interaction, whereas the miRNA-mRNA interaction existing in at least a half of cells or tissues are defined as a conservative miRNA-mRNA interaction. Given the identified sample-specific miRNA regulatory networks, we can obtain dynamic and conservative miRNA-mRNA interactions which form dynamic and conservative miRNA regulatory networks, respectively.

### Hub miRNA identification

Hub miRNAs are defined as highly connected miRNAs in a dynamic and conservative miRNA regulatory network. In this work, we use the cumulative probability of Poisson distribution to evaluate whether a miRNA is a hub:

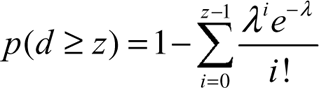

where *λ* = *np*, 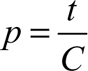, *n* is the number of genes (including miRNAs and mRNAs), *t* is the number of miRNA-mRNA interactions in a dynamic or conservative miRNA regulatory network, and *C* is the number of all possible miRNA-mRNA interaction pairs. Smaller *p*-value of a miRNA shows that the miRNA is more likely to be a hub. Here, the cutoff of the *p*-value is set to 0.05.

### Validation analysis

To understand whether the identified dynamic and conservative hub miRNAs are closely associated with CML and BRCA, we get a list of CML-related and BRCA-related miRNAs from RNADisease v4.0^75^. As a result, we have obtained 203 CML-related and 1493 BRCA-related miRNAs (**Supplementary Data 9**). If a dynamic or conservative hub miRNA is overlapped with the list of CML-related or BRCA-related miRNAs, the dynamic or conservative hub miRNA is regarded as a CML-related or BRCA-related miRNA. **Comparison metrics**

We use two metrics (accuracy and efficiency) to compare different instances of Scan using different network inference methods, and compare Scan with other methods for inferring sample-specific miRNA regulation. For accuracy, the ground truth of miRNA-mRNA interactions are acquired from miRTarBase v9.0^76^ and TarBase v8.0^77^ for validation. If a method has a larger percentage of validated miRNA-mRNA interactions, the method will have higher accuracy. For efficiency, we compare the runtime of different methods in the same bulk or single-cell RNA-sequencing data. If a method takes less runtime in the same bulk or single-cell RNA-sequencing data, the method will have better efficiency.

In terms of accuracy and efficiency, we use an overall rank score^78^ to evaluate the performance of each method. For the method *i*, the overall rank score *ors_i_* is calculated as:

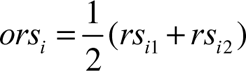

where *rs_i_*_1_ and *rs_i_* _2_ denote the rank scores of method *i* in terms of accuracy and efficiency, respectively.

A method with higher accuracy or better efficiency will obtain a larger rank score. A method with a larger overall rank score is regarded as a better or practical method.

### Method execution

Each execution of Scan and CSmiR on bulk or single-cell RNA-sequencing data is performed in a separate task. Each task is allocated 32 CPU cores of Intel(R) Xeon(R) Platinum 8375C CPU at 2.90 GHz, and one R session is opened for each task. The network inference methods of Scan with runtime more than 10 days or with memory usage more than 256 GB are discarded.

## Data availability

All accession codes, unique identifiers, and web links for publicly available datasets are described in the paper. All data supporting the findings of the current study are listed in Supplementary Data and our GitHub website (https://github.com/zhangjunpeng411/Scan/).

## Code availability

Scan is released under the GPL-3.0 License, and is available at https://github.com/zhangjunpeng411/Scan/.

## Acknowledgements

This work has been supported by the National Natural Science Foundation of China (61963001), the Yunnan Fundamental Research Projects (202001AT070024, 202101BA070001-221), and the ARC DECRA (DE200100200).

## Author contributions

J.Z. and T.D.L. conceived the idea of this work. L.L. and J.L. refined the idea. J.Z. designed and performed the experiments. X.W., C.Z. and Y.L. participated in the design of the study and performed the statistical analysis. J.Z., L.L., T.D.L. and J.L. drafted the manuscript. All authors revised the manuscript. All authors read and approved the final manuscript.

## Competing interests

The authors declare no competing interests.

## Notes

https://github.com/zhangjunpeng411/Scan

### Competing Interest Statement

The authors have declared no competing interest.

