## Supplementary File 1 for "Scanning sample-specific miRNA regulation from bulk and single-cell RNA-sequencing data"

Supplementary Figures

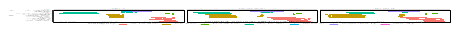

**Supplementary Fig. 1 Cell-specific miRNA regulation using linear interpolation strategy (Scan.interp) in the K562 dataset. a** Cell-specific miRNA regulatory networks with and without using priori information. **b** Power law degree distribution of cell-specific regulatory networks with and without using priori information. **c** Similarity of cell-specific regulatory networks with and without using priori information. “None” denotes no priori information of miRNA targets used, “TargetScan” is the priori information of miRNA targets used from TargetScan database, and “ENCORI” represents the priori information of miRNA targets used from ENCORI database.

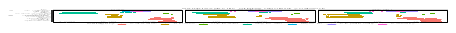

**Supplementary Fig. 2 Cell-specific miRNA regulation using statistical perturbation strategy (Scan.perturb) in the K562 dataset. a** Cell-specific miRNA regulatory networks with and without using priori information. **b** Power law degree distribution of cell-specific regulatory networks with and without using priori information. **c** Similarity of cell-specific regulatory networks with and without using priori information. “None” denotes no priori information of miRNA targets used, “TargetScan” is the priori information of miRNA targets used from TargetScan database, and “ENCORI” represents the priori information of miRNA targets used from ENCORI database.

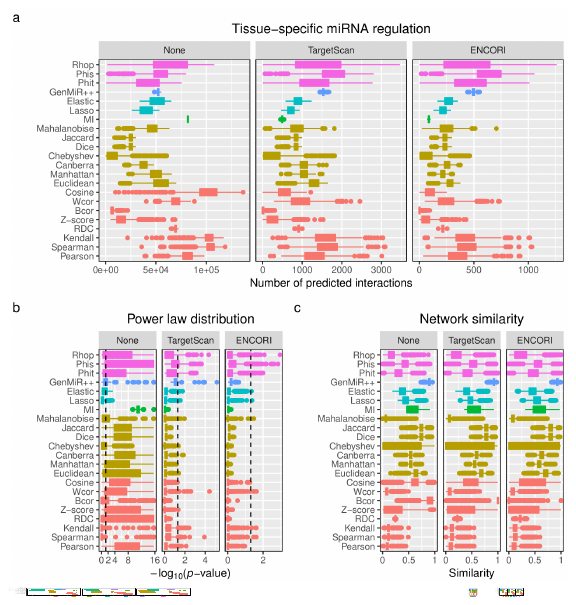

**Supplementary Fig. 3 Tissue-specific miRNA regulation using linear interpolation strategy (Scan.interp) in the BRCA dataset. a** Tissue-specific miRNA regulatory networks with and without using priori information. **b** Power law degree distribution of tissue-specific regulatory networks with and without using priori information. **c** Similarity of tissue-specific regulatory networks with and without using priori information. “None” denotes no priori information of miRNA targets used, “TargetScan” is the priori information of miRNA targets used from TargetScan database, and “ENCORI” represents the priori information of miRNA targets used from ENCORI database.

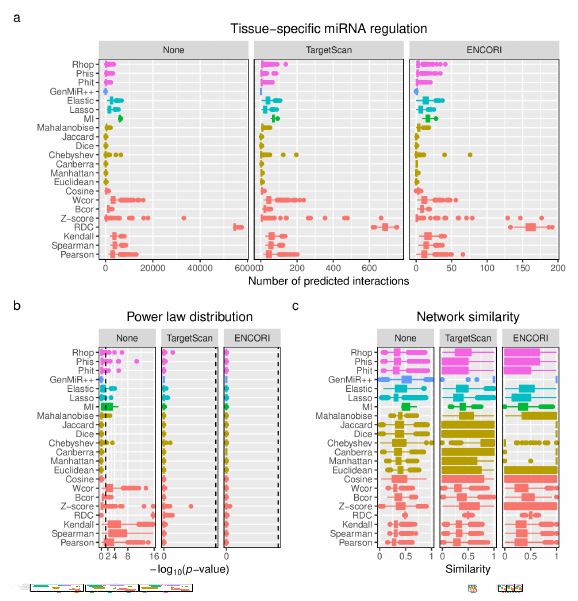

**Supplementary Fig. 4 Tissue-specific miRNA regulation using statistical perturbation strategy (Scan.perturb) in the BRCA dataset. a** Tissue-specific miRNA regulatory networks with and without using priori information. **b** Power law degree distribution of tissue-specific regulatory networks with and without using priori information. **c** Similarity of tissue-specific regulatory networks with and without using priori information. “None” denotes no priori information of miRNA targets used, “TargetScan” is the priori information of miRNA targets used from TargetScan database, and “ENCORI” represents the priori information of miRNA targets used from ENCORI database.

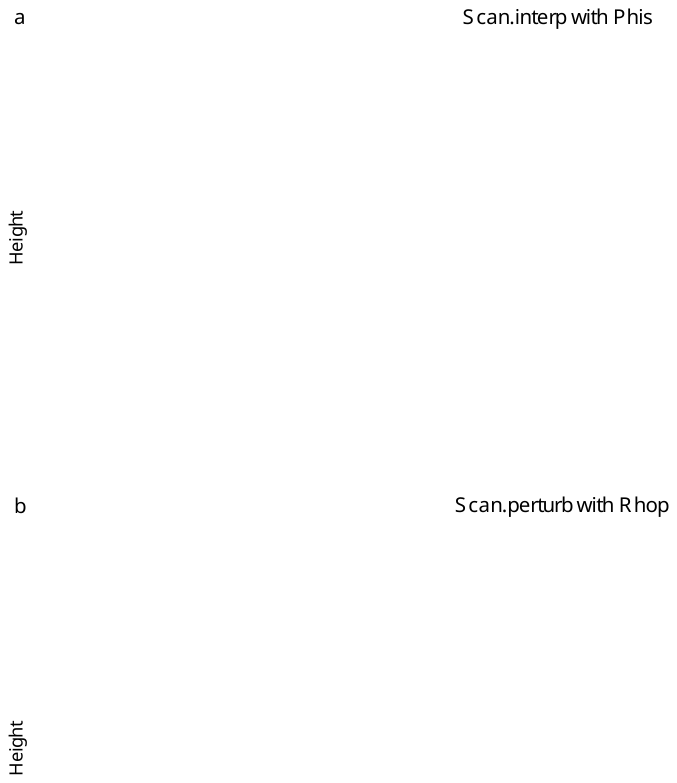

**Supplementary Fig. 5 Clusters of BRCA tissues. a** Clusters of BRCA tissues, and Scan.interp with Phis is used to identify tissue-specific miRNA regulatory networks. **b** Clusters of BRCA tissues, and Scan.perturb with Rhop is used to infer tissue-specific miRNA regulatory networks. Each color denotes a cluster.
